# High-throughput Monoclonal Antibody Screening from Immunized Rabbits via Droplet Microfluidics

**DOI:** 10.1101/2025.03.27.645851

**Authors:** Johnson Q. Cui, Ruyuan Song, Weihong Song, Ouyang Li, Xin Yuan, Hongbo Zhou, Lu Zhang, Shuhuai Yao

**Affiliations:** Department of Mechanical and Aerospace Engineering, The Hong Kong University of Science and Technology, Clear Water Bay, Kowloon, Hong Kong 999077, China; Department of Chemical and Biological Engineering, The Hong Kong University of Science and Technology, Clear Water Bay, Kowloon, Hong Kong 999077, China; DPBIO, Inc., 324 S Diamond Bar Blvd, Unit Num 717 Diamond Bar, CA 91765, United States

**Keywords:** monoclonal antibody, microfluidics, droplet sorting, high throughput screening, magnetic negative selection

## Abstract

The discovery of monoclonal antibodies (mAbs) is critical to advancing therapeutics, diagnostics, and biomedical research. While mouse-derived mAbs dominate current applications, their limitations—short serum half-life, human immunogenicity, and restricted recognition of human-specific antigens—underscore the need for alternative sources. Rabbit-derived mAbs have been gaining significant traction with their superior antigen-binding affinity, broader epitope diversity, and higher yield potential. However, the absence of well-defined surface markers on rabbit B cells has hindered efficient enrichment strategies, limiting the exploration of this valuable antibody repertoire. In this study, we present an integrated workflow that combines magnetic negative selection with high-throughput droplet microfluidics to overcome these barriers. By optimizing a pan B cell enrichment protocol using a tailored antibody cocktail, we achieved a notable boost in IgG secretion and B cell enrichment. Through two complementary droplet-encapsulated assays using particle aggregation for soluble antigens and reporter cells for membrane-bound antigens, we identified target cells capable of secreting high-affinity IgGs. Subsequent sequencing, *in vitro* antibody production and characterization confirmed the high affinity rate of the discovered antibodies, outperforming rates previously reported. The use of droplet microfluidics streamlines the analysis of rabbit IgG repertoires, providing a robust platform for rabbit single B cell antibody discovery with promising applications in precision medicines and diagnostics.

## Introduction

Monoclonal antibodies (mAbs) have revolutionized therapeutics, diagnostics, and biomedical research by enabling precise targeting of antigens with high specificity^1–3^. In therapeutics, mAbs underpin the principles of precision medicine, offering tailored treatments based on individual patient needs and disease characteristics. In diagnostic applications, mAbs are instrumental in accurately and swiftly detecting biomarkers, pathogens, and disease-related molecules, facilitating early disease detection and monitoring. In research settings, mAbs are indispensable tools for probing protein functions, elucidating cellular mechanisms, and unravelling disease pathways with high specificity. Since the first FDA approval of a therapeutic mAb (anti-CD3) in 1986, which was aimed at preventing organ transplant rejection, over 100 mAbs had received FDA approval by 2021^4^.

Conventional mouse-derived mAbs, while supported by established protocols, suffer from limitations including short serum half-life^5–7^, immunogenicity risks (*e.g*., human anti-mouse antibody reactions)^8–10^, and weak responses to human-specific antigens^11–13^, restricting their therapeutic and diagnostic utility. In contrast, rabbit mAbs offer superior affinity, specificity, and broader epitope recognition on human antigens^14, 15^, facilitating the discovery of cross-reactive antibodies and providing higher yields due to rabbits’ larger size. Thus, efficient, high-throughput methods for assessing rabbit mAbs are increasingly valuable.

Traditional mAb discovery relies on *in vivo* immunization followed by immortalization (e.g., hybridoma technology^16, 17)^ or immune cell libraries in display methods (e.g., phage^18^ or yeast^19^), and screening. Though reliable, hybridoma methods are slow and inefficient due to low fusion rates, species limitations, and bias toward plasmablasts, while display techniques demand labor-intensive library construction and multiple screening rounds, historically struggling to retain natural heavy chain (V_H_) and light chain (V_L_) pairing^20, 21^, thus reducing diversity. These drawbacks have spurred the development of alternative, more efficient mAb discovery technologies.

Recent advances in phenotypic screening of antibody-secreting cells (ASCs, plasma cells and plasmablasts) have shifted from low-throughput ELISpot assays to miniaturized and automated systems. For instance, methods utilizing on-chip valves^22^, hydrodynamic cell traps^23^, microwells^24^, or micropillar arrays^25^ enable higher throughput of ~10^2^-10^3^ cells. However, their broader adoption is still hindered by manual handling, limited well/array capacity, and difficulties in retrieving positive clones. The commercial platform Beacon™ utilizes the principle of light-induced dielectrophoresis, also referred to as OptoElectro Positioning (OEP), to precisely manipulate ASCs within thousands of nanopens^26^. Even higher throughput can be obtained by droplet microfluidic-based methods, which can generate millions of highly monodisperse droplets at high speed, and each droplet functions as an individual reaction vessel that can be sorted^27^, injected^28, 29^, or merged^30^ based on specific needs. More importantly, the encapsulation of cells and their secretions within droplets enables a direct linkage between the cells’ phenotype and genotype. This miniaturization of reaction volumes (which are typically several orders of magnitudes smaller than wells) allows secreted biomolecules to reach detectable concentrations in shorter time, empowering various functional assays with high sensitivity, efficiency, and versatility^31–33^. Droplet microfluidics techniques enable streamlined isolation of antibody-secreting cells (ASCs) from mice at the single-cell level based on antigen-binding activity, followed by paired V_H_-V_L_ sequencing, bioinformatics analysis, and *in vitro* antibody production and characterization^34–36^. Furthermore, a method was developed by combining an antibody capture hydrogel with antigen bait in flow cytometry to quickly and efficiently identify mAbs from mouse and human ASCs^37^. To simultaneously quantify the functional activities and production rate of both cell-surface and secreted proteins from individual cells, a Förster resonance electron transfer (FRET)-based detection assay encapsulated in droplets can differentiate functional hybridoma cells in less than 30 min^38^.

Despite these exciting advancements, most existing research focused on the screening of mouse-derived mAbs, whereas the exploration of rabbit-derived mAbs has been significantly underdeveloped. One significant obstacle stems from the absence of universal surface markers for rabbit B cells, compounded by the lack of commercially accessible kits for enriching rabbit B cells. Currently, fluorescence activated cell sorting (FACS) is typically used to isolate antigen-specific memory B cells, but this method does not facilitate the screening of ASCs, which typically produce antibodies with superior affinity and higher quantity compared to those from memory B cells, as they express few or no immunoglobulins (Igs) on their surface, leading to the loss of potentially valuable specific antibodies. Furthermore, the staining steps involved in this method can adversely affect cell viability, complicating subsequent sequence recovery. While platforms like Beacon™ employ rabbit memory B cell activation, this approach is constrained by factors such as long activation cycles, low efficiency, poor sample-sample consistency, and reduced diversity. Therefore, to accelerate mAb mining from rabbits, there is an urgent need for a robust, stable, and efficient antibody screening method capable of identifying rabbit B cells that secret antibodies of interest.

In this work, we established an integrated platform for rapid rabbit mAb discovery by combining a customized rabbit pan B cell enrichment protocol with high-throughput droplet microfluidics. By optimizing an antibody cocktail with a diverse set of antibodies, magnetic negative selection was used to enrich B cells from rabbit spleens, achieving a sevenfold enrichment in IgG secretion (validated by ELISpot) and a ninefold increase in B cell enrichment (confirmed by single-cell sequencing). To screen antigen-specific IgG^+^ B cells, we encapsulated two complementary assays within water-in-oil droplets: (1) a particle aggregation-based assay for soluble antigens and (2) a reporter cell-based assay for cell surface antigens. Antigen-specific droplets were identified through multi-channel spatial fluorescence profiling, followed by sorting and subsequent single-cell lysis and sequencing, enabling recovery of cognate V_H_-V_L_ gene pairs for in-vitro antibody production. The antibodies’ binding affinity was assessed via ELISA or flow cytometry, with 90% (9/10) binding soluble ovalbumin (OVA) and 70% (7/10) binding cell surface antigen hCD82. By leveraging droplet microfluidics, this workflow enables high-throughput mining of the rabbit IgG repertoire within ~2 weeks from spleen harvest to antibody validation, offering a scalable and efficient platform for antibody discovery with broad applications in therapeutic, and diagnostic, and research applications.

## Results and Discussion

### Streamlined Workflow of mAb Discovery from Rabbit

Leveraging the high-throughput nature of droplet microfluidics, a streamlined workflow for rapid discovery of mAbs from immunized rabbits is developed and illustrated in Figure 1a. To address the diversity of natural antigen forms, which exist as both soluble entities (*e.g*., shed proteins in blood) and cell-associated structures (*e.g.*, on cancer cells), and to broaden the applicability of our platform, we integrated a particle aggregation-based assay for a soluble antigen and a reporter cell-based assay for a cell surface antigen, aiming for more comprehensive and versatile mAb mining.

**Figure 1.**
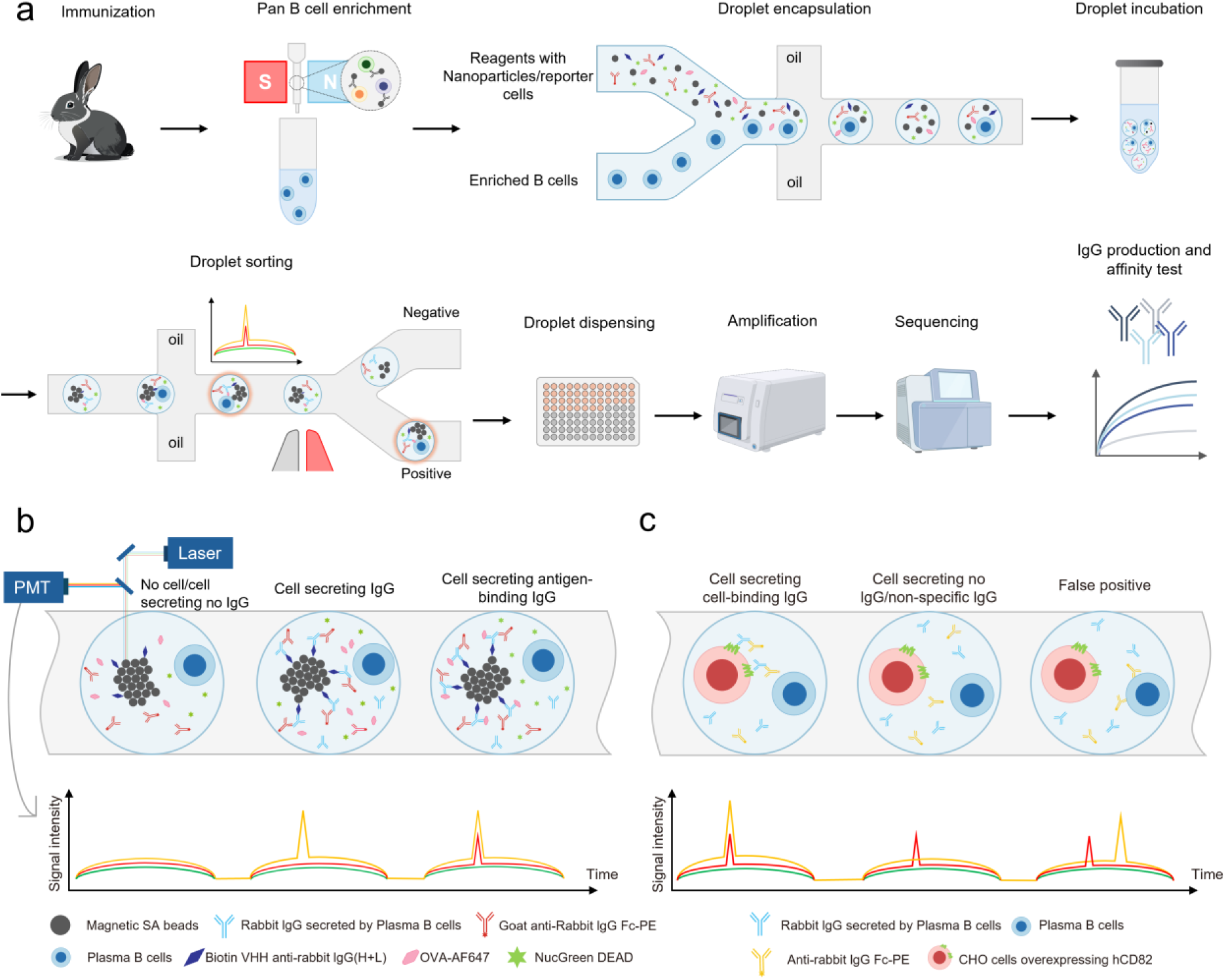
Process flow and mechanism of rabbit mAb discovery via droplet microfluidics. a. Process flow of mAb discovery from rabbits immunized with soluble antigens OVA, or 293T cells overexpressing hCD82, which consists of 9 steps including rabbit immunization, pan B cell enrichment, droplet encapsulation, droplet incubation, droplet sorting, droplet dispensing, sequencing, and IgG production and affinity test. b, c. Working mechanism of the droplet screening assay for soluble antigens OVA (b) and cell membrane antigen hCD82 (c). Only the droplets that contain cells secreting antibodies with binding affinity to the corresponding antigens exhibit colocalized fluorescence signal peaks in yellow and red channels.

In our mAb discovery workflow, New Zealand rabbits were first immunized intraperitoneally with either a soluble antigen (OVA) or 293T cells overexpressing hCD82 to elicit an immune response. Following immunization, splenocytes were harvested, and B-cell lineages were enriched using an optimized magnetic negative selection method. This enrichment step enhanced the efficiency of subsequent single-cell encapsulation by removing non-B cells. The enriched B cells were then collected and prepared for encapsulation in picoliter-sized droplets using a dual aqueous-phase microfluidic droplet generation chip. The resulting droplets were incubated at 37°C for an hour to allow IgG secretion and detection via the in-drop assays.

To detect cells secreting IgGs with binding affinity to the soluble antigen OVA, a particle aggregation-based assay was constructed to screen target cells based on the colocalized fluorescence signals on the particle aggregation. During droplet generation, the aqueous phase 1 is the enriched cell suspension diluted to 4.5 million cells/ml to minimize the droplets containing more than one cell dictated by Poisson distribution, resulting in the mean cell number per droplet being 0.3. The aqueous phase 2 contains paramagnetic nanoparticles coated with biotin VHH anti-rabbit IgG (H+L), Goat anti-Rabbit IgG Fc Fragment specific-PE, OVA protein conjugated with Axela Fluro 647, and the dead cell staining reagent NucGreen dead. As indicated in Figure 1b, within droplets that contain cells secreting target antibodies, the antibodies were first captured onto the paramagnetic colloidal nanoparticles coated with biotin VHH anti-rabbit IgG (H+L), which spontaneously formed a particle aggregate to reduce the surface energy^39^. In cases where the secreted antibody was an IgG, the Goat anti-Rabbit IgG Fc Fragment specific-PE within the droplet relocated onto the particle aggregate and emitted yellow fluorescence (EX: 561 nm, EM: 575 nm). Furthermore, if the IgG had binding affinity for OVA labeled with AF647, the antigens also relocated to the aggregate and emitted red fluorescence (EX: 635 nm, EM: 647 nm). While within droplets that contained no cells or non-IgG secreting B cells, none of the mentioned signal was triggered. Within droplets that contained dead cells, green fluorescence was triggered by NucGreen dead (EX: 488 nm, EM: 521 nm). Using this in-drop fluorescent sandwich immunoassay, only the droplets that containing cells that secreted target IgGs exhibited colocalization of dual fluorescence at the particle aggregate, but not the remainder. Furthermore, the incorporation of the particle aggregate not only mitigates the low frequency of droplets containing both a single cell and a single microbead, a common issue dictated by double Poisson statistics in other approaches, but also it increases binding efficiency due to its larger surface area and strengthens fluorescence signal intensity through aggregation.

To detect cells secreting IgGs with binding affinity to the cell surface antigen hCD82, a reporter cell-based assay was constructed to screen positive droplets based on colocalization of fluorescence signals on the reporter cell surface. For the droplet generation, instead of nanoparticles and soluble antigens, CHO cells (Chinese hamster ovary cells) overexpressing hCD82 were stained with CellTrace Far Red and added in aqueous phase 2, which were co-compartmentalized with enriched pan B cells in droplets, with a mean of 1 reporter cell and 0.3 B cell per droplet, maximizing droplets with single B cell while ensuring the presence of reporter cells for the binding affinity assay. As indicated in Figure 1c, similarly, only the droplets emitting colocalized IgG signal (yellow fluorescence) and reporter cell signal (red signal) at the reporter cell surface contained IgGs with target binding affinity, but not the remainder.

A 220 µL total volume of the two aqueous phases, mixed at a 1:1 ratio, was emulsified in the droplet generator using an inert fluorinated carrier oil, yielding approximately 3.3 million droplets with a diameter of ~50 µm. After sufficient droplet incubation, the droplets were subsequently reinjected and sorted using a microfluidic sorting chip according to their spatial fluorescence characteristics. Each droplet was subjected to scanning and excitation through superimposed laser lines, with fluorescent signals recorded via photomultiplier tubes (PMT). Droplets with dual fluorescence colocalization at the particle aggregate or reporter cell surface indicated desired IgG secretion, which were sorted and dispensed into a 96-well plate via fluorescence-activated dielectrophoretic force. The cells collected by the 96-well plate were then sequenced, and cognate *V_H_* − *V_L_* pairs were identified. Antibodies were subsequently expressed via transfection in 293T cells, and the affinity of the expressed antibodies were further assessed by ELISA for solution antigen OVA and flow cytometry analysis for cell surface antigen hCD82.

### B Cell Enrichment

B cell enrichment is a critical step in achieving high throughput mAb mining, removing irrelevant cells (*e.g.*, T cells, macrophages, and other non-B cells) increases the likelihood of encapsulating a target B cell, enhances the efficiency of identifying rare ASCs, and minimizes contaminants that could lead to false positives in binding assays or interference with PCR amplification of antibody genes, ensuring cleaner data and more reliable identification of mAbs.

Following the immunization of rabbits and the harvest of spleen, splenocytes were collected by grinding and 600-1200 million cells were obtained per spleen. To optimize the negative selection method to enrich B cells, series of antibody cocktails were prepared by mixing biotinylated anti-rabbit CD4 antibody, anti-rabbit CD8 antibody, anti-rabbit T lymphocytes antibody, anti-rabbit IgM antibody, anti-human CD11b antibody, anti-human CD123 antibody, anti-human CD11c antibody at various concentrations, which were specific to surface markers on non-B cells, e.g., T cells, monocytes, dendritic cells, etc. Moreover, these antibodies are biotinylated, enabling the tagged non-B cells to be captured later using streptavidin beads. The cell mixture was then passed through an LS column, which retained the bead-bound non-B cells using magnetic field, while the unlabeled cells (enriched B cells) flowed through. After assessing the enrichment results via single-cell sequencing (Supplementary. Figure S1), we selected the enrichment protocol with the best performance, which achieved a ninefold enrichment of B cells (Figure. 2a) and a sevenfold increase in IgG secretion measured by ELISpot (Figure. 2b). Meanwhile, the cell viability remained high from 90% before enrichment to 94% after enrichment (Figure. 2c) and the cell number decreased by 90% (Figure. 2d), signifying significant depletion of non-target cells. The B enrichment strategy was also tested by enriching B cells in peripheral blood mononuclear cells (PBMCs) of rabbits (Supplementary. Figure S2), validating its versatility in different samples. Furthermore, the enrichment procedure significantly reduced cell aggregation upon loading in the microfluidic chip, facilitating single cell encapsulation in droplets (Supplementary. Figure S3).

**Figure 2.**
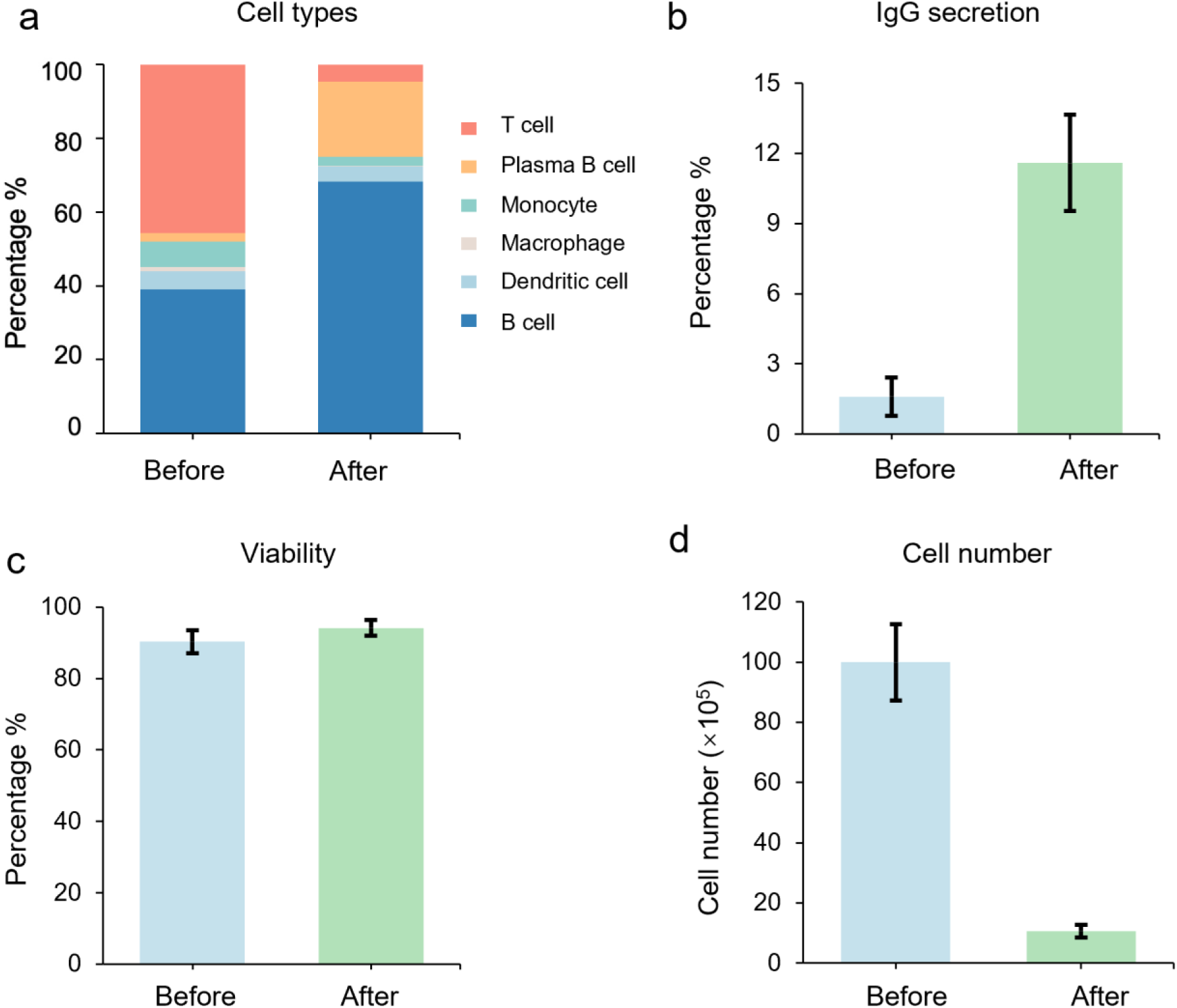
Pan B cell enrichment by magnetic negative depletion. a. Cell types in the spleen sample before and after enrichment measured by single cell sequencing, a ninefold enrichment of B cells was achieved. b. IgG secretion of the spleen sample before and after enrichment measured by ELISpot, a sevenfold increase in IgG secretion ratio was observed. c. Cell viability remained high (>90%) before and after cell enrichment. d. Cell number drastically decreased by 90% after cell enrichment.

### Droplet Encapsulation, Incubation, and Sorting

After B cell enrichment, the cells were encapsulated in droplets with bioreagents using a flow focusing microfluidic droplet generation chip, which were then collected and incubated at 37 °C for an hour to enable sufficient antibody secretion and detection via the in-drop fluorescent sandwich immunoassay. A microscopy view of the droplets in droplet generation chip and droplet sorting chip is shown in Figure. 3a (particle aggregation-based assay), and Figure. 3b (reporter cell-based assay). After incubation, droplets were observed under a fluorescence microscope, and representative positive droplets images in different fluorescence channels were shown in Figure. 3c (particle aggregation-based assay) and Figure. 3d (reporter cell-based assay).

**Figure 3.**
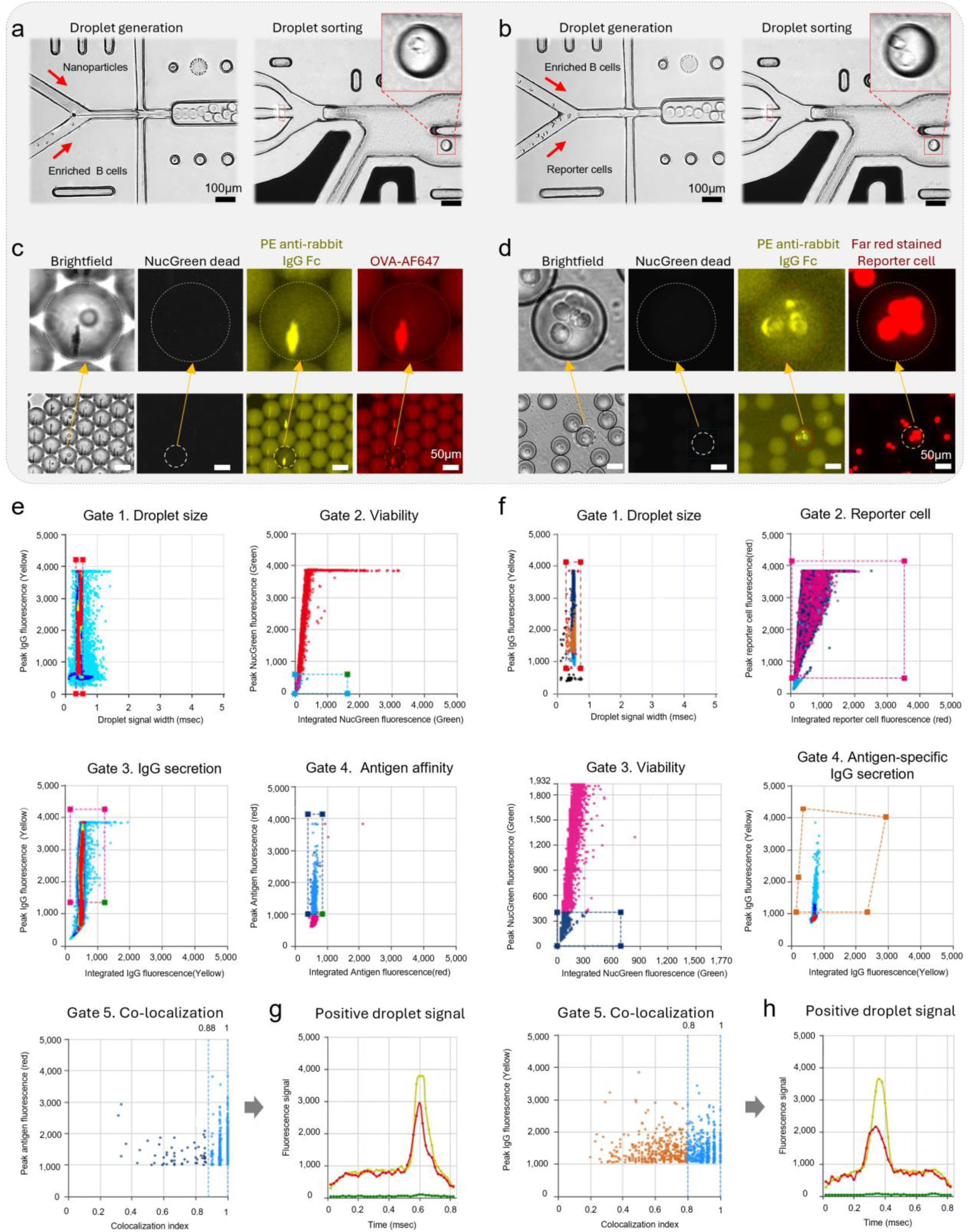
Droplet generation, sorting, and gating strategy. a, b. Microscopy images of droplet generation and droplet sorting for particle aggregation-based assay(a) and reporter cell-based assay(b) respectively, scale bar is 100 µm. c, d. Droplet images in 4 optical channels (brightfield, green, yellow, and red fluorescence channel) after incubation, enlarged images show the target droplet containing cell secreting IgG with binding affinity to OVA (c), or hCD82 (d). e. The droplet gating strategy for screening primary B cells producing IgG with affinity to OVA. 5 gates were applied sequentially to sort droplets based on following parameters or states, 1. Droplet size, 2. Cell viability, 3. IgG secretion, 4. Affinity to OVA, 5. Colocalization indexes of signals. f. The droplet gating strategy for screening primary B cells producing IgG with affinity to membrane antigen hCD82. 5 gates were applied sequentially to sort droplets based on following parameters or states, 1. Droplet size, 2. Reporter cell signal, 3. Cell viability, 4. Antigen specific IgG signal, 5. Colocalization indexes of signals. g, h Representative fluorescence signal images of positive droplets from both methods obtained by PMT. All fluorescence values are in arbitrary units (a.u.).

To select cells with the desired IgG secretion, the incubated droplets were reinjected into a microfluidic sorting chip for fluorescence-activated dielectrophoretic sorting based on their multi-channel spatial fluorescence characteristics. As illustrated in Figure. 3e, f, a multi-step sequential gating strategy was deployed to set the criteria for droplet sorting for detecting soluble antigen binding and cell surface antigen binding respectively, in these scatter plots, the droplets within the region enclosed by a dashed outline were selected while the rest were considered negative and discarded. In Figure. 3e, firstly, the droplets sample was gated to screen droplets of desired sizes with diameters ranging from 48 to 52 um and eliminate coalesced droplets and smaller broken droplets (Gate 1). Secondly, the droplets were gated to exclude those containing dead cells that emitted green fluorescence levels surpassing the established threshold (Gate 2). Thirdly, the droplets’ yellow and red fluorescence were analysed and gated to screen droplets with significant fluorescence signal levels in both channels (Gate 3, 4). Finally, to quantify the colocalization of the fluorescence signals in yellow and red channels on the particle aggregate, we defined a colocalization index, c, as c = 1-*t_interval_*/*t_passage_*, with *t_interval_* representing the time interval in between the peaks in the yellow and red fluorescence channels, and *t_passage_* representing the passage time of the whole droplet. Thus, c is naturally constrained within the range of 0 to 1, with a value of 1 signifying the complete colocalization of the two peaks in the yellow and red fluorescence channels. To determine the colocalization of the two fluorescence signals, a threshold of 0.88 was chosen and droplets exhibiting higher colocalization indexes were considered as positive droplets and sorted (Gate 5).

In Figure. 3f, a similar gating strategy was employed for assessing droplet size (Gate 1), cell viability (Gate 2), reporter cell signal (Gate 3), antigen-specific IgG signal (Gate 4), and the colocalization of the fluorescence signal for IgG signal and reporter cell signal (Gate 5). The droplets with colocalization indexes higher than 0.8 were sorted. Representative fluorescence signal images of positive droplets from both methods obtained by PMT are shown in Figure. 3g, h respectively. Apart from sorting sample droplets containing enriched B cells, optically barcoded negative and positive control groups were also differentiated and analysed in the sorting process (Supplementary. Figure. S4, S5). The droplets were sorted at a rate of ~500 Hz, with the entire process requiring approximately 2 hours to identify positive events from a pool of ~3.3 million droplet samples.

According to the gating and sorting results, in the particle aggregation-based assay, the droplet containing cells with IgG secretion made up 10.28 % of all the compartmentalized cells from the enriched cell sample. And among the cells that secreted IgG, 2.1 % showed specific bindings to the antigen OVA. For the reporter cell-based assay, 0.81 % of the compartmented cells exhibited specific bindings to the cell membrane hCD82, validating our platform’s ability to screen rare functional events from vast droplet libraries. The positive droplets were sorted then dispensed individually into a 96-well plate containing storage buffer, the plate with sorted droplets was sealed with aluminum foil seals for downstream processes.

### Antibody Production and Binding Affinity Test

After droplet generation, incubation, sorting and dispensing, the plates dispensed with positive droplets were treated to disrupt the emulsion and release the cells. Subsequently, lysis buffer was added to facilitate cell lysis, followed by reverse transcription and cDNA amplification. The V_H_ and V_L_ genes were amplified using nested PCR. The products from the second PCR round were purified, sequenced via Sanger sequencing, and analyzed bioinformatically with the International ImMunoGeneTics Information System (IMGT) database to annotate variable region gene segments. Subsequently, the V_H_ and V_L_ genes were constructed as linear expression cassettes, which were then fused into full-length immunoglobulin (Ig) V_H_ and V_L_ cassettes by overlapping PCR, ensuring precise segment integration.

The evolutionary relationships among heavy chain (IGHV) and light chain (IGKV) immunoglobulin gene sequences were represented by circular phylogenetic trees illustrated in Figure. 4a, b. The phylogenetic trees exhibit a radial layout where branches radiate from a central node, and the length and branching patterns reflect the genetic distance or similarity between sequences. The trees are color-coded to distinguish different gene families or subgroups, with labels indicating specific gene identifiers, offering a visual representation of the phylogenetic relationships among IGHV and IGKV gene families respectively, highlighting their diversity and evolutionary connections. 10 assemblies were constructed from the particle aggregation-based assay and the reporter cell-based assay each. For antibody production, the PCR products of the linear Ig expression cassettes were purified and co-transfected into 293T cells, which were cultured and incubated. The supernatants were collected and the IgGs were purified by affinity chromatography. The produced IgGs’ affinity was assessed by ELISA for the soluble antigen OVA, and by flow cytometry for cell surface antigen hCD82. For the ELISA test, the antibodies were serially diluted 3-fold, ranging from 0.01 nM to below 100 nM, together with a blank negative control group and a positive control group, the absorbance signals were measured and plotted in Figure. 4c. As the result indicated, 9 out of 10 antibodies showed binding affinity towards OVA. For the flow cytometry test, precultured hCD82-CHO cells were harvested and incubated with 20nM of purified antibodies, the cells were washed and incubated with 50 nM of mouse anti-rabbit PE-conjugated secondary antibody at 4 °C for 30 minutes, and the subsequent flow cytometry results were shown in Figure. 4d, where the x-axis represents fluorescence values, and the y-axis represents cell counts. As the results indicated, 7 out of 10 antibodies showed binding affinity towards hCD82 and exhibited similar fluorescence profiles to the positive control group, which is significantly higher than previously reported 14%^36^. This improvement may stem from our mAb discovery workflow, where rabbits were immunized with 293T cells overexpressing hCD82, while CHO cells expressing hCD82 were used as reporter cells in the droplet assay to ensure IgG specificity. In contrast, previous works relied on a single cell line for both immunization and screening, potentially increasing the selection of non-specific IgGs targeting non-target antigens. The relatively lower affinity rate of the reporter cell-based method compared to particle aggregation-based method could be due to its lower efficiency in the immunization strategy, *i.e.*, purified soluble antigen versus transfected reporter cell. Additionally, in the reporter cell assays, non-specific interactions such as antibody sticking to reporter cell components could trigger false positive events. In general, our affinity assessments substantiated that our mAb discovery strategy via droplet microfluidics effectively examined B cells from rabbits, with a predominant number of the discovered mAbs showing significant affinity towards the antigens.

**Figure 4.**
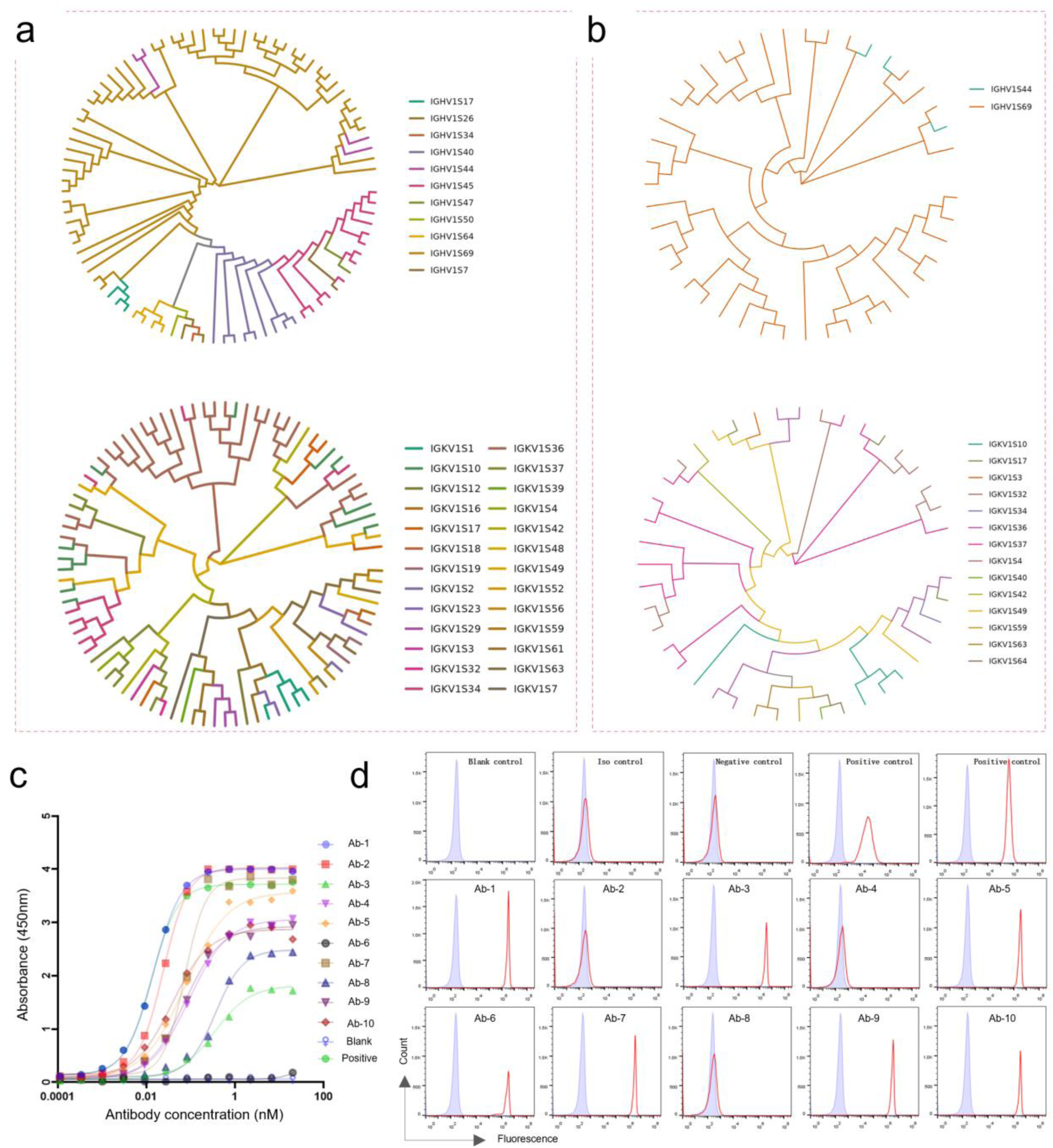
Sequencing, production and characterization of IgGs expressed by the sorted cells. In rabbits immunized with OVA (a) and hCD82 (b), the diversity of V-gene families in paired V_H_ and V_L_ sequences was evaluated through phylogenetic trees, which were built from full-length V-region amino acid sequences and color-coded by their closest germline V-gene family. c. Affinity test of the 10 antibodies obtained from particle-aggregation assay with positive and negative control group via ELISA, the results showed that 9 antibodies demonstrated significant binding affinity towards antigen OVA. d. Affinity test of the 10 antibodies obtained from reporter cell-based assay with positive and negative control group via flow cytometry, the results showed that 7 antibodies demonstrated binding affinity toward cell surface antigen hCD82.

### Conclusion

The development of advanced methods for mAb discovery holds profound implications for therapeutics, diagnostics, and biomedical research. While mouse-derived mAbs remain predominant, their limitations including short half-life, human immunogenicity, and poor recognition of human-specific antigens constrain their utility for clinical use. Rabbit mAbs, in contrast, offer distinct advantages: superior antigen-binding affinity, broader epitope diversity (particularly against human antigens), and higher yield potential. However, the exploration of rabbit mAbs has been hindered by the lack of efficient B cell enrichment techniques, largely due to the absence of well-defined surface markers on rabbit B cells.

In this work, we address this challenge by integrating a pan B cell enrichment strategy with high-throughput droplet microfluidics to establish a rapid, streamlined workflow for mining the rabbit IgG repertoire. By optimizing a magnetic negative selection protocol using a tailored antibody cocktail, we achieved a sevenfold increase in IgG secretion (via ELISpot) and a ninefold enrichment of B cells (via single-cell sequencing). To screen antigen-specific IgG+ B cells, we engineered two complementary droplet-encapsulated assays: (1) a particle aggregation-based assay for soluble antigens (OVA) and (2) a reporter cell-based assay for membrane-bound antigens (hCD82). Target cells were identified based on their muti-channel spatial florescence characteristics, followed by sorting, lysis, and sequencing to recover paired heavy-chain and light-chain gene sequences. These sequences were expressed by in-vitro production of antibodies through transfection in 293T cells, and the affinity of the produced antibodies was assessed by ELISA or flow cytometry, revealing that 90% (9/10) bound antigen OVA and 70% (7/10) recognized cell surface hCD82, a significantly higher affinity rate than previously reported for antibodies targeting cell surface antigens. By combining phenotypic screening with single-cell genomics, our droplet microfluidics-driven workflow enables comprehensive interrogation of rabbit IgG diversity within ~2 weeks from spleen harvest to antibody validation, overcoming traditional bottlenecks in B cell enrichment and screening. This platform not only accelerates the discovery of high-affinity rabbit mAbs but also expands their potential applications in precision therapeutics, biomarker detection, and mechanistic research.

## Materials and Methods

### Immunization of Rabbits

Immunization with OVA: New Zealand rabbits were immunized intraperitoneally with 400 μg ovalbumin(OVA; Sigma) in complete Freund’s adjuvant (Sigma) mixed 1:1 with 0.9% (wt/vol) NaCl (Versylene Fresenius) for primary immunization (day 0), Secondary, tertiary, and fourth immunizations (days 14, 28, and 42) used OVA emulsified in incomplete Freund’s adjuvant (Sigma) mixed 1:1 with 0.9% NaCl, while the final boost (day 52) used OVA in 0.9% NaCl alone. Immunization with hCD82: New Zealand rabbits were immunized intravenously with ~10^7^ 293T cells overexpressing hCD82 in PBS for primary immunization (day 0), followed by intraperitoneal immunizations every 3 weeks (days 21, 42, and 63). The final boost (day 77) involved intravenous injection of 100 μg hCD82-hFc.

### Cell Line Construction

Two reporter cell lines, 293T overexpressing hCD82 (for immunization) and CHO overexpressing hCD82 (for screening), were generated through transfection and antibiotic selection. A hCD82/KAI-1 ORF cDNA expression plasmid (Sino Biological) was separately transfected into 293T and CHO cells using Lipo3000 (Thermo Fisher). Transfected cells were selected with hygromycin, and high-hCD82-expression cells were sorted using a Beckman CytoFLEX SRT.

### Extraction of Cells from Spleen and B-cell Enrichment

Splenocytes were harvested from the spleens of rabbits on day 55 of the immunization schedule for soluble antigens (3 days post-boost). Spleens were mechanically ground to collect cells, yielding an average of 600–1200×10^6^ cells per spleen after red blood cell lysis. Non-B cells were depleted using biotinylated anti-rabbit antibodies against CD4, CD8, T lymphocytes (all Bio-Rad), and IgM (BD Biosciences), plus anti-human antibodies against CD11b, CD123, and CD11c (all Thermo Fisher), followed by streptavidin beads (Miltenyi) and passage through an LS column (Miltenyi). The unlabeled cells, representing enriched B cells, showed an approximately eight-fold increase in IgG-secreting cells (plasmablasts and plasma cells) by ELISpot.

### ELISpot for Cell Enrichment Assessment

IgG-secreting cells before and after pan B-cell enrichment were enumerated using ELISpot. A 96-well filtration plate (Millipore, MSIPS4510) was activated with 35% ethanol for 30 seconds and washed five times with 1× PBS. Wells were coated overnight with 100 μL of 1 mg/mL goat anti-rabbit IgG Fc secondary antibody (Sino Biological, SSA017). After washing five times with 1× PBS, wells were blocked with cell culture medium (X-VIVO, supplemented with 10% low-IgG FBS) for 30 minutes at 37°C. Subsequently, 5,000 cells from pre- and post-enrichment samples were seeded in the wells with cell culture medium and incubated for 16 hours. IgG secretion was detected by incubating with 1 mg/mL biotinylated goat anti-rabbit IgG (H+L) antibody (Jackson ImmunoResearch) for 2 hours, followed by 1:1000 diluted streptavidin-ALP (Mabtech, 3310-10) for 30 minutes, and developed with BCIP/NBT-plus (Mabtech, 3650-10). Images of each well were captured under a microscope.

### Single-cell RNA-seq Library Construction

Single-cell RNA sequencing (scRNA-seq) libraries were constructed using the Galaxy Single Cell Analysis System (DPBIO,SCI100) according to the manufacturer’s instructions^40^. Cell suspensions were filtered through a 70-μm strainer, washed twice with cold Dulbecco’s PBS (D-PBS), and resuspended in D-PBS with 0.04% BSA. Cell viability and concentration were assessed using a Luna-FL™ cell counter (Logos Biosystems). Approximately 20,000 cells (at 1,000 cells/μL, >90% viability) were loaded into each channel of a single-cell isolation chip, targeting ~10,000 cells per channel. Following encapsulation, cells were lysed in droplets, and mRNA was captured and barcoded via reverse transcription. The emulsion was then broken, and cDNA was amplified with cycle settings optimized for different tissue types. Amplification products were analyzed on an Agilent 4200 instrument (Agilent Technologies), followed by fragmentation, end repair, A-tailing, index adapter ligation, and library amplification per the kit’s user guide. Libraries were sequenced on an MGI T7 platform (MGI, China) at a depth of at least 20,000 reads per cell.

### Single-Cell RNA-seq Analysis

Single-cell RNA sequencing (scRNA-seq) data underwent quality control and preprocessing using cutadapt (v 4.1) to remove sequencing adapters and trim low-quality 3′ nucleotides^41^. Reads shorter than 50 nucleotides after trimming were discarded. Quality-filtered reads were aligned to the rabbit reference genome (Oryctolagus cuniculus v2.0) using STARsolo^42^ (v 2.7.11b). and cells were identified using the EmptyDrops algorithm in CellRanger-compatible mode to generate a cell/UMI count matrix^43^. Gene annotations were then converted to human homologs with the biomaRt package^44^. Data analysis was performed in Seurat, following standard workflows: log-normalization, identification of highly variable genes, data scaling and centering, and dimensionality reduction. To address batch effects, multiple samples were integrated using Seurat’s canonical correlation analysis (CCA) method, leveraging sample-specific principal component analysis (PCA) results.

### Aqueous Phases for Cell Compartmentalization

#### Particle aggregation-based droplet assays

Two aqueous phases were prepared for cell encapsulation and affinity assays in droplets. For the sample group, Aqueous Phase 1 consisted of cell suspensions (4.5 × 10⁶ cells/mL) prepared at 4°C by resuspending cells in DMEM/F12 supplemented with 0.1% Pluronic F68 (Life Technologies), 25 mM HEPES (pH 7.4; Life Technologies), 5% Knockout Serum Replacement (Life Technologies), and 1% Pen/Strep (Thermo Fisher), targeting a λ (mean number of cells per droplet) of ~0.3 for B cells in soluble antigen-binding assays. Aqueous Phase 2 comprised paramagnetic nanoparticles (Streptavidin Plus, 300 nm; Beaver) coated with biotinylated VHH anti-rabbit IgG (H+L) (AlpVHH) and resuspended in a working solution of 80 nM goat anti-rabbit IgG Fc-specific PE (Jackson ImmunoResearch), 100 nM OVA-AF647, and 6% (v/v) NucGreen Dead (resulting in final droplet concentrations of 40 nM anti-rabbit IgG Fc-PE and 50 nM antigen). For the negative control, Aqueous Phase 1 replaced B cells with 50 nM rabbit IgG isotype (GenScript) with 250 nM Alexa Fluor 405 carboxylic acid for droplet barcoding, while Aqueous Phase 2 remained identical to the sample group,

#### Reporter cell-based droplet assays

For the sample group, Aqueous Phase 2 contained CHO-hCD82 reporter cells stained with 50 μM CellTrace Far Red (Thermo Fisher) and resuspended in a working solution with 80 nM goat anti-rabbit IgG Fc-specific PE (Jackson ImmunoResearch) and 6% (v/v) NucGreen Dead, while Aqueous Phase 1 matched the sample group from the soluble antigen-binding assays. For negative and positive controls, Aqueous Phase 1 contained either rabbit anti-hCD82 antibody (Invitrogen) with 1,000 nM Alexa Fluor 405 carboxylic acid (positive control) or rabbit IgG isotype (GenScript) with 250 nM Alexa Fluor 405 carboxylic acid, both spiked into supplemented DMEM/F12 (as above) to a final concentration of 50 nM; Aqueous Phase 2 was unchanged from the sample group.

### Droplet Generation, Collection, Incubation, and Imaging

After preparation, the two aqueous phases (220 μL total volume) were mixed at a 1:1 ratio, co-flowed, and partitioned into droplets using a flow-focusing microfluidic chip (DPBIO, CRF0041). The continuous phase was fluorinated droplet generation oil. The operating pressures of the droplet generation chips were adjusted to produce monodisperse droplets (50 ± 5 μm diameter) at approximately 5,500 Hz (Aqueous Phase 1: 3.5 psi; Aqueous Phase 2: 3.5 psi; oil phase: 10 psi). Droplets were collected into a 1.5-mL DNA LoBind tube (Eppendorf) immediately after generation, sealed, and incubated at 37°C for 1 hour to allow antibody secretion and reagent interactions. Post-incubation, droplets were imaged under a fluorescence microscope to assess fluorescence on nanoparticle aggregation or the reporter cell surface.

### Droplet Sorting, Gating strategy and Dispensing

Droplet fluorescence analysis and sorting were performed on a CytoSpark Droplet Sorting System (DPBIO, CSP3400, MRPT9A), where a fixed-focus laser line (wavelengths of 405 nm, 488 nm, 561 nm, or 635 nm) was oriented perpendicular to the sorting channel and fluorescence was analyzed using a photomultiplier tube. During sorting, the gating threshold for the antibody-binding signal (yellow fluorescence) was set at 1.5× the fluorescence of blank droplets, while the antigen-binding signal (red fluorescence) threshold was adjusted based on the negative control group to ensure non-specific signals from the negative control were below 0.1%. Desired droplets were sorted via dielectrophoretic force with the following parameters: Q_emulsion = 3.5 psi, Q_spacer oil = 3.2 psi, Q_sheath oil = 4.2 psi, sorting pulse frequency = 10 kHz, sorting pulse duration = 1,200 μs, and sorting voltage = 1.3 kV. Sorted single cells were dispensed into a 96-well plate, each well containing 8 μL of storage buffer (0.25 μL of RNase inhibitor [40 U/μL; ABclonal], 10mM DTT, and 7.75 μL of nuclease-free water). After sorting and dispensing, plates were sealed with aluminum foil until further use.

### Single Cell cDNA Synthesis and Amplification

The cDNA synthesis and amplification were performed using a Single-Cell cDNA Synthesis and Amplification Kit (DPBIO, RRPF17B) according to the manufacturer’s instructions. Plates containing dispensed droplets were retrieved from storage, and each well was supplemented with 2.5 μL of emulsion recovery agent. Subsequently, 4 μL of lysis buffer was added to each well, followed by incubation at 72°C for 3 minutes in a thermal cycler. The lysed droplet contents were then mixed with 8 μL of reverse transcription (RT) buffer and transferred to a PCR plate. RT was conducted in a thermal cycler with the following conditions: 42°C for 90 minutes, 10 cycles of 50°C for 2 minutes and 42°C for 2 minutes, 85°C for 5 minutes, and a hold at 4°C. Next, 25 μL of PCR mix containing 0.3 μL of 100 μM ISPCR primer was added, followed by nuclease-free water to a final volume of 50 μL per well. cDNA amplification was performed under these conditions: 98°C for 3 minutes, 22 cycles of 98°C for 20 s, 65°C for 30 s, and 72°C for 4 minutes, then 72°C for 5 minutes, and a hold at 4°C. The amplified cDNA was either used directly or stored at −20°C.

### BCR Amplification and Sanger Sequencing

The VH and VL genes were amplified via two-round nested PCR. PCR conditions for both rounds were as follows: 95°C for 3 minutes, 30 cycles of 95°C for 15 s, 60°C for 15 s, and 72°C for 90 s, followed by 72°C for 5 minutes, and held at 4°C. First-round forward and reverse primers, specific to VH and Vκ leader sequences and heavy and kappa constant regions^45^. Second-round primers for VH (variable heavy chain) and JH (joining heavy chain) regions are listed in Supplementary Table 1. Second-round PCR products were purified using AMPure XP beads at a 0.8:1 (v/v) ratio and eluted in 40 μL of nuclease-free water, then subjected to Sanger sequencing. Sequencing primers are provided in Supplementary Table 1. The resulting sequences were analyzed with the IMGT Information System (https://www.imgt.org/) to identify variable region gene segments.

### Assembly of Ig VH and VL gene Linear Expression Cassettes

Linear expression cassettes for Ig VH and VL genes were assembled as previously described^46^. Each cassette included essential transcriptional and translational elements: a CMV promoter, Ig leader sequences, constant regions of human IgG1 heavy chain (IgHC) or kappa light chain (IgLC), a poly(A) tail, and substitutable VH or VL genes. The CMV, IgHC, and IgLC fragments were *de novo* synthesized and cloned into pUC57 plasmids (General Biol). For cassette assembly, these fragments were amplified from the plasmids via PCR using primers listed in Supplementary Table 2. Full-length Ig heavy- and light-chain expression cassettes were constructed by overlapping PCR. The heavy-chain cassette combined CMV and IgHC fragments with 2 μL of purified second-round VH PCR products, while the kappa light-chain cassette used CMV and IgLC fragments with 2 μL of purified second-round VL PCR products. Each 50-μL PCR reaction contained 25 μL of KAPA HiFi HotStart ReadyMix (Roche), 10 μM CMV_Forward primer, 10 μM IgHC_Reverse or IgLC_Reverse primer, and water. Amplification conditions were 95°C for 3 minutes, 30 cycles of 95°C for 15 s, 60°C for 15 s, and 72°C for 2 minutes, followed by 72°C for 5 minutes, and held at 4°C.

### Antibody Expression and Purification

PCR products of the linear Ig expression cassettes were purified using AMPure XP beads (Beckman Coulter) at a 0.8:1 (v/v) ratio and eluted in 40 μL of nuclease-free water. Twenty-four hours prior, 293T cells were seeded in 6-well plates at a density of 300,000–500,000 cells per well. The following day, purified PCR products of the paired Ig heavy- and light-chain gene expression cassettes were co-transfected into 293T cells using Lipo293™ transfection reagent (Beyotime, C0521). Six to eight hours post-transfection, the medium was replaced with fresh culture medium supplemented with 2% (v/v) fetal calf serum (FCS), and cells were incubated at 37°C in a 5% CO₂ incubator for 72 hours. Supernatants were then collected, clarified by centrifugation at 500g for 10 minutes at room temperature, and stored at 4°C. IgG was subsequently purified using affinity chromatography on a Protein G gravity column (Protein G Sepharose 4 fast flow column, GE Healthcare).

### ELISA for Affinity Test

The antigen (1 μg/mL OVA) was coated onto a microtiter plate and incubated at 4°C overnight. The following day, the antigen solution was discarded, and the plate was blocked with 2% (w/v) BSA in PBS. Purified IgG from supernatants (initial concentration: 100 nM) was serially diluted threefold over 11 steps in PBS. After incubation at 37°C for 0.5 h, the antibody solution was removed, and the plate was washed five times with 300 μL of PBST (PBS with 0.05% Tween-20). An HRP-conjugated secondary antibody (0.15 μg/mL; Thermo Fisher) was added, incubated at 37°C for 0.5 h, then discarded, followed by five washes with 300 μL of PBST. Color was developed using o-phenylenediamine chromogenic solution, the reaction was stopped with ELISA Stop Solution (Solarbio), and the OD₄₅₀ was measured using a microplate spectrophotometer.

### Flow Cytometry for Affinity Test

Pre-cultured hCD82-CHO cells were harvested and seeded into 96-well plates at 1 × 10^6^ cells per well. Cells were washed three times with cold Dulbecco’s PBS (DPBS; 4°C), resuspended in 100 μL of cold DPBS (Thermo Fisher), and incubated with 20 nM purified antibodies from supernatants at 4°C for 30 min. Subsequently, cells were washed three times with cold DPBS, resuspended in 100 μL of cold DPBS, and incubated with 50 nM phycoerythrin (PE)-conjugated mouse anti-rabbit secondary antibody (Jackson ImmunoResearch) at 4°C for 30 min. After three additional washes with cold DPBS, cells were resuspended in 100 μL of DPBS and analyzed by flow cytometry on a BD FACSymphony™ A1 instrument (BD Biosciences).

## Supporting information

Additional B cell enrichment data, control groups for droplet assays, and sequences used.

## Author Contributions

All authors have given approval to the final version of the manuscript.

## Notes

*R.S., W.S., O.L., X.Y., and H.Z.* performed the research as part of their employment at DPBIO. The other authors declare no competing financial interest.

## Acknowledgements

We thank DPBIO, Inc for providing the resources and support necessary to conduct this research.

